# Synchronisation of apical constriction and cell cycle progression is a conserved behaviour of pseudostratified neuroepithelia informed by their tissue geometry

**DOI:** 10.1101/2022.06.15.496231

**Authors:** Ioakeim Ampartzidis, Christoforos Efstathiou, Francesco Paonessa, Elliott M Thompson, Tyler Wilson, Conor J McCann, Nicholas DE Greene, Andrew Copp, Frederick J Livesey, Nicola Elvassore, Giovanni Giuseppe Giobbe, Paolo De Coppi, Eirini Maniou, Gabriel L Galea

## Abstract

Neuroepithelial cells balance tissue growth requirement with the morphogenetic imperative of closing the neural tube. They apically constrict to generate mechanical forces which elevate the neural folds, but are thought to apically dilate during mitosis. However, we previously reported that mitotic neuroepithelial cells in the mouse posterior neuropore have smaller apical surfaces than non-mitotic cells. Here, we document progressive apical enrichment of non-muscle myosin-II in mitotic, but not non-mitotic, neuroepithelial cells with smaller apical areas. Live-imaging of the chick posterior neuropore confirms apical constriction synchronised with mitosis, reaching maximal constriction by anaphase, before division and re-dilation. Mitotic apical constriction amplitude is significantly greater than interphase constrictions. To investigate conservation in humans, we characterised early stages of iPSC differentiation through dual SMAD-inhibition to robustly produce pseudostratified neuroepithelia with apically enriched actomyosin. These cultured neuroepithelial cells achieve an equivalent apical area to those in mouse embryos. iPSC-derived neuroepithelial cells have large apical areas in G2 which constrict in M phase and retain this constriction in G1/S. Given that this differentiation method produces anterior neural identities, we studied the anterior neuroepithelium of the elevating mouse mid-brain neural tube. Instead of constricting, mid-brain mitotic neuroepithelial cells have larger apical areas than interphase cells. Tissue geometry differs between the apically convex early midbrain and flat posterior neuropore. Culturing human neuroepithelia on equivalently convex surfaces prevents mitotic apical constriction. Thus, neuroepithelial cells undergo high-amplitude apical constriction synchronised with cell cycle progression but the timing of their constriction if influenced by tissue geometry.

## Introduction

Embryonic cells must balance physiological requirements for growth with energy-intensive force generating behaviours necessary to form functional organs during morphogenesis. Closure of the neural tube is a clinically relevant paradigm of morphogenesis. It requires bending of the neural plate, which elevates paired neural folds that meet at the dorsal midline, forming closure points from where “zippering” initiates to progressively close the open neuropores (Nikolopoulou et al., 2017). Failure to close the neural tube causes neural tube defects in an average of 1:1,000 births globally (Zaganjor et al., 2016). Incomplete closure of the anterior neuropores causes fatal anencephaly whereas failure to close the posterior neuropore (PNP) causes spina bifida. Both cranial and spinal neural tube closure require neuroepithelial cells to undergo apical constriction (Butler et al., 2019, Galea et al., 2021, Lesko et al., 2021, Kowalczyk et al., 2021).

Apical constriction is a conserved force-generating behaviour whereby actomyosin recruitment shrinks epithelial the apical domain of cells. It has been intensively studied in *Drosophila, Xenopus* and nematodes, identifying common and distinct constriction mechanisms (Martin and Goldstein, 2014). Some cells preferentially recruit actomyosin to the apicomedial cell surface, assembling radial F-actin filaments which guide constriction towards the middle of the cell’s apex (Coravos and Martin, 2016). Other cell types preferentially recruit myosin to the cell cortex, serially and often directionally shrinking cell junctions (Nishimura et al., 2012). Much of the contractile machinery is common to both constriction modalities. Both require non-muscle myosin motor proteins such as myosin-IIb, activated through phosphorylation by kinases including Rho-associated kinase (ROCK). Phosphorylated myosins localise to the cell apical surface where they bind F-actin scaffolds and generate contractile forces through ATP-dependent power strokes. Both apicomedial and cell junction actomyosin can contribute to apical shrinkage. In the stratified neuroepithelium of *Xenopus*, apicomedial F-actin accumulation is associated with apical shrinkage in the presumptive brain, whereas both apicomedial and cortical accumulation can correlate with apical shrinkage in the spinal region (Baldwin et al., 2022).

We have documented both apicomedial and cortical myosin-IIb localisation in the pseudostratified neuroepithelium of the mouse PNP during spinal neurulation (Galea et al., 2021). Apicomedial myosin-IIb localisation correlates with dense microtubule networks in both interphase and mitotic cells, in which they are apical to, and distinct from, the mitotic spindle microtubules (Galea et al., 2021). Cortical localisation of myosin heavy chain (MHC)-IIb is diminished in non-constricting neuroepithelial cells whose neighbours lack the planar cell polarity (PCP) co-receptor VANGL2. ROCK is a downstream effector of PCP signalling known to enhance neuroepithelial apical constriction in multiple species, including the mouse and chick (Butler et al., 2019, Nishimura et al., 2012). Its pharmacological inhibition globally reduces neuroepithelial tension, enlarges apical surfaces and diminishes apical phospho-myosin-IIb localisation in the mouse PNP (Butler et al., 2019, Escuin et al., 2015). We previously reported that ROCK’s effects on apical constriction are related to cell cycle progression. As neuroepithelial cells proliferate in the presence of ROCK inhibition, the proportion of cells with dilated apical surfaces increases and the PNP progressively widens (Butler et al., 2019). This demonstrates the importance of coordination between apical constriction and cell proliferation. Excessive neuroepithelial proliferation has repeatedly been associated with failure of neural tube closure in mouse genetic models (Lardelli et al., 1996, Anderson et al., 2016, Parchem et al., 2015, Badouel et al., 2015).

Cell cycle progression in the highly proliferative neuroepithelium is unusual because these cells are pseudostratified in amniotes. Consequently, their nucleus displaces along the apical-basal axis due to interkinetic nuclear migration (IKNM) as cells progress through the cell cycle. Mitoses are predictably localised apically, followed by stochastic basal descent such that S phase nuclei are predominantly basal (Leung et al., 2011). As cells pass through G2 phase their nuclei rapidly ascend apically. G2 nuclear ascent is an active process which can be achieved by different cellular mechanisms in tissues with different geometries. For example, in the flat neuroepithelium of the zebrafish hindbrain it requires ROCK-dependent myosin activation, whereas in the apically convex developing retina it requires formin-mediated sub-nuclear F-actin accumulation (Yanakieva et al., 2019a). These displacements require nuclear deformation, which is opposed by nuclear envelope proteins such as Lamin A/C (Swift et al., 2013). Embryonic neuroepithelia characteristically lack A-type Lamin expression (Jung et al., 2012, Takamori et al., 2018, Yanakieva et al., 2019a) and over-expression of Lamin A diminishes their nuclear motility during IKNM (Yanakieva et al., 2019b).

IKNM is believed to reverse neuroepithelial apical constriction by causing apical dilation during mitosis (Guerrero et al., 2019). The dogma that mitosis causes apical expansion is strongly supported by findings in *Drosophila* showing reversal of apical constriction and loss of adherens junctions during mitosis (Ko et al., 2020, Aguilar-Aragon et al., 2020). However, it does not hold true in the mouse PNP neuroepithelium: we previously reported that neuroepithelial cells labelled with proliferating cell markers such as phospho-histone H3 (pHH3) unexpectedly have smaller apical areas than interphase cells (Butler et al., 2019). We therefore propose that synchronisation of apical constriction with cell cycle progression is a neuroepithelial behaviour which reconciles physiological growth requirements with the necessity of closing the neural tube.

As a recently described cell behaviour, very little is known about the regulation of neuroepithelial mitotic apical constriction. Here, we set out to document its relation to apical myosin recruitment, timing during mitotic progression, and conservation between species. We document that synchronisation of apical constriction with cell cycle progression is a conserved property, including in human cells differentiated from induced pluripotent stem cells (iPSCs). However, whereas flat neuroepithelia undergo mitotic apical constriction, mitotic dilation predominates in apically convex neuroepithelia of the early mouse midbrain and human cells cultured on convex surfaces.

## Results

### Mouse posterior neuropore neuroepithelial cells accumulate apical myosin IIb

Neuroepithelial cells have tortuous shapes with numerous sub-apical protrusions (Kasioulis et al., 2022), variable nuclear positioning along the apicobasal axis and often unpredictable regional widening or narrowing of the cell body (Figure 1A). Here, we quantify apical surface properties related to each cell’s body or nucleus using 3D visualisation aided with basolateral markers (scribbled or adherens junction markers). Individual cells can be traced to relate deep nuclei to the cell’s apical surface as previously described (Butler et al., 2019). Apically located mitotic nuclei do not protrude beyond the PNP epithelial surface, rather sitting below a constricted cell apex (Figure 1A) frequently decorated with ROCK (Figure 1B).

**Figure 1:**
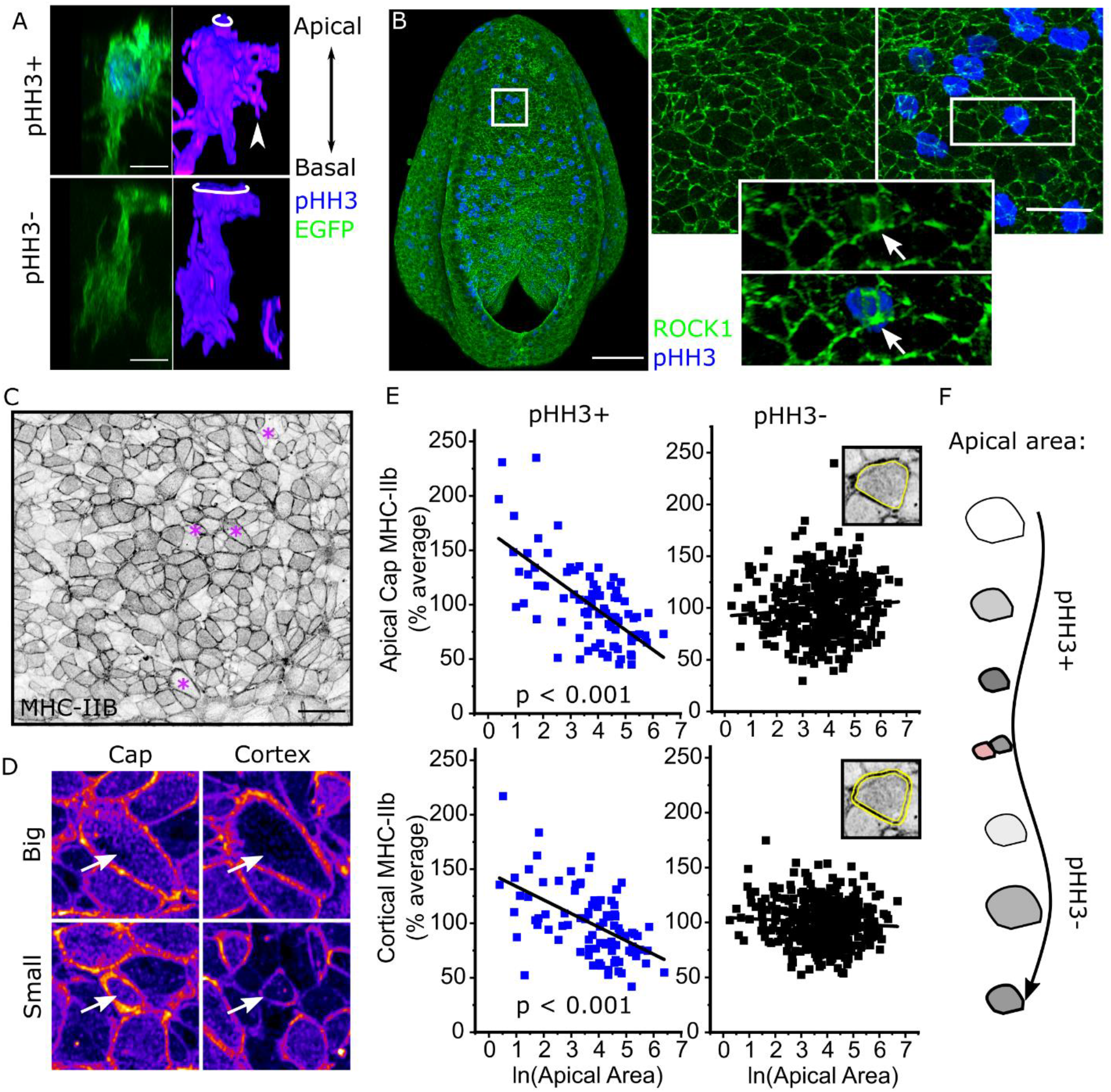
Mitotic neuroepithelial cells in the mouse PNP have constricted apical surfaces with enriched myosin. **A.** 3D reconstructions showing the shape of illustrative EGFP mosaically-labelled neuroepithelial cells positive or negative for pHH3. EGFP lineage-tracing was induced with Nkx1.2^CreERT2^. White rings indicate the apical surface, the arrowhead indicates a sub-apical protrusion. Scale = 10 μm. **B.** Wholemount immunofluorescence showing neuroepithelial apical localisation of ROCK1 and pHH3. Arrow indicates apical surface of a mitotic cell in a 3D reconstruction. Scale = 100 μm (full PNP) and 25 μm (inset). **C-D.** Surface-subtracted wholemount immunofluorescence showing neuroepithelial apical localisation of MHC-IIB. Scale = 20 μm. * in **C** indicate cells shown in **D** (fire LUT in which faint staining is in blue and bright staining in yellow) to illustrate variability in cap/cortical MHC-IIb irrespective of apical area. **E.** Correlations between apical area and MHC-IIb intensity. Each embryo’s MHC-IIb values are normalised to its average staining intensity to correct for inter-individual differences. Yellow lines in insets show the cap versus cortex regions analysed. pHH3+ = 90 cell and pHH3- = 401 cells from 8 embryos with 17-19 somites. **F.** Schematic illustrating the proposed association between apical area and apical MHC-IIb. pHH3+ cells progressively accumulate MHC-IIb (grey) as they constrict their apical surface. Following division, pHH3- cells can either constrict or dilate.

MHC-IIb is localised in apicomedial pools (“cap”) and/or circumferentially at cell-cell junctions (“cortical”, Figure 1C,D). Mitotically rounded cells continue to localise MHC-IIb around their apical cap and/or cortex similarly to non-mitotic cells, as well as in their sub-apical cell cortex (Supplementary Figure 1). Individual cells’ apicomedial cap and cortical MHC-IIb intensities are correlated in both mitotic and non-mitotic cells (mitotic R^2^ = 0.51, p < 0.001; non-mitotic R^2^ = 0.48, p < 0.001). Neither apical cap nor cortical MHC-IIb intensity is correlated with the apical area of interphase (pHH3-) cells (Figure 1E). In contrast, both myosin pools are inversely correlated with apical area in mitotic cells (pHH3+, Figure 1E), suggesting that progressive accumulation of apical myosin predictably shrinks their apical surface. This association may not be evident in static analyses of interphase cells due to the pulsatile nature of apical constriction (Christodoulou and Skourides, 2015, Galea et al., 2021). Cells with equivalent apical areas may have been either constricting or dilating at the point of embryo fixation (Figure 1F).

### Mitotic apical constriction is the highest magnitude constriction in the chick spinal neuroepithelium

The chick spinal neuroepithelium is more amenable to dynamic analyses requiring long-term live imaging than the mouse. Species differences are notable: average apical areas are smaller in the chick than mouse PNP neuroepithelium (Figure 2A-B). Nonetheless, mitotic neuroepithelial cells have smaller apical areas than their corresponding interphase cells in both chick and mouse embryos (Figure 2B), confirming evolutionary conservation of mitotic apical constriction. Apical areas of mouse mitotic neuroepithelial cells were on average 40.7% smaller than non-mitotic cells (pHH3^+^ 20.5 μm^2^ versus pHH3^-^ 34.5 μm^2^) and those of chick embryos decreased by 33% (pHH3^+^ 13.5 μm^2^ versus pHH3^-^ 19.2 μm^2^).

**Figure 2:**
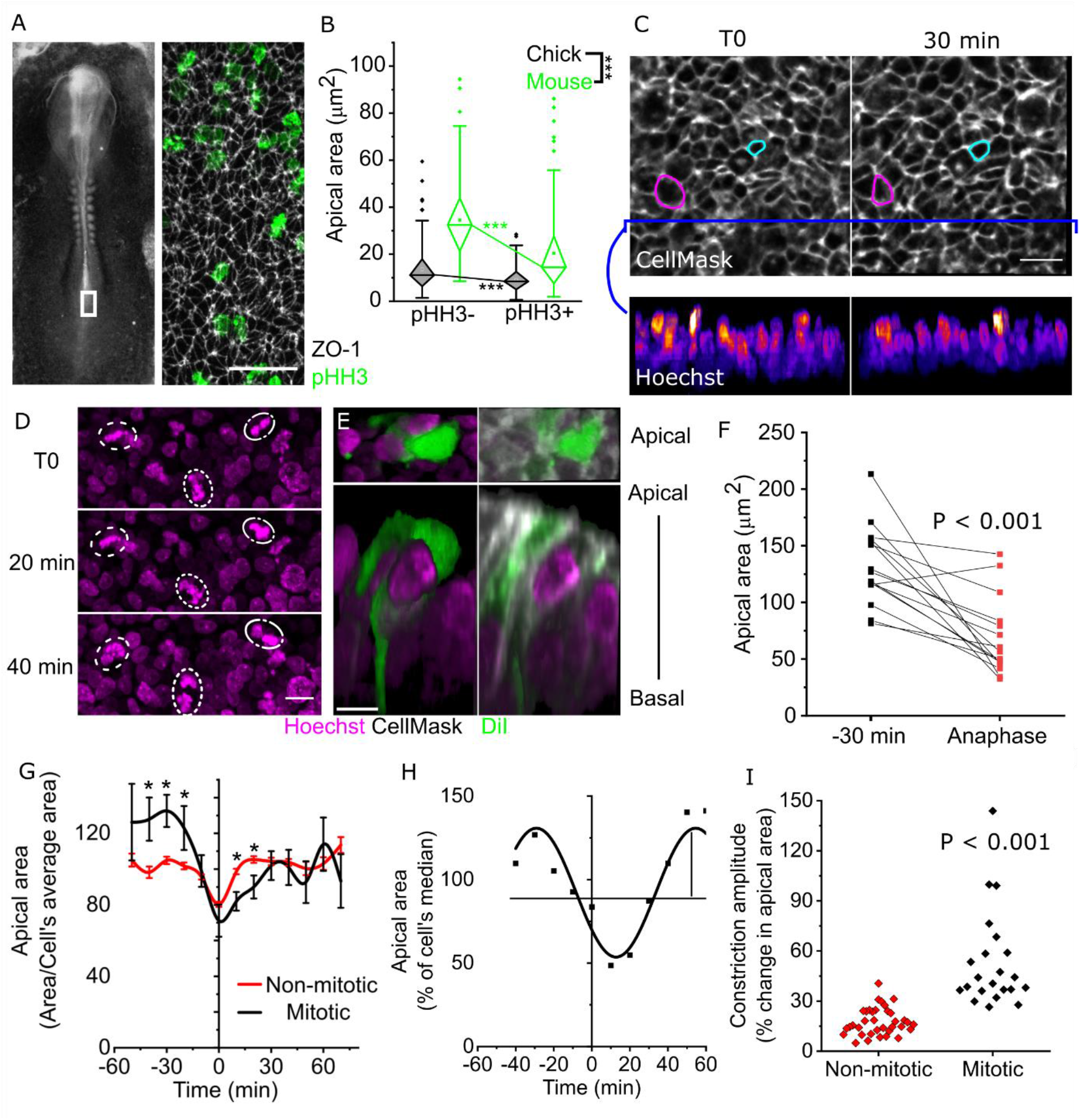
Neuroepithelial apical constriction occurs between G2 and anaphase in the chick spinal neuroepithelium. **A.** Illustrative brightfield image showing the approximate region (white box) visualised following wholemount immunolabelling for ZO-1 and pHH3. Scale = 50 μm. **B.** Apical area quantification of pHH3+ and pHH3- cells in mouse or chick PNPs. ***p < 0.001 by two-way ANOVA with Bonferroni post-hoc. Chick = 143 pHH3+ and 186 pHH3- cells from 5 embryos. Mouse = 108 pHH3+ and 108 pHH3- cells from 4 embryos. **C.** Snapshots of live-imaged chick neuroepithelium showing the apical surface in grey LUT and optically resliced nuclei in Fire LUT showing changes over 30 mins. Magenta = apical constriction, cyan = dilation. Scale = 20 μm. **D.** Live-images snapshots showing Hoechst-labelled nuclei or mitotic chromatin. Dotted lines indicate the same mitotic chromatin in different time points. Scale = 10 μm **E**. Apical and rostro-caudal 3D reconstructions of live-imaged cells labelled with mosaic DiI as positional landmarks, CellMask used to analyse the apical surface (as in **C**), and Hoechst to identify cells which undergo mitosis. Scale = 10 μm. **F.** Apical area quantification of mitotic cells with an anaphase nuclear morphology and 30 mins prior. Points represent individual cells. P value from paired t-test. **G.** Quantification of live-imaged apical area normalised to each cell’s average apical area at all timepoints analysed. Cell traces are aligned such that T0 indicates anaphase nuclear morphology in mitotic cells and the minimum observed apical area of non-mitotic cells. Lines represent the mean ± SEM, numbers of cells at each time point vary (T0 = 54 non-mitotic and 29 mitotic cells from 4 independent embryos). * p < 0.05 by repeated measures ANOVA. **H.** Illustrative sine curve fitting to the apical area values of a single live-imaged cell. Arrow indicates the curve amplitude. **I.** Comparison of apical area curve amplitude between mitotic and non-mitotic cells. P value by T-test.

The chick neuroepithelium can be live-imaged with vital cell dyes through a window in the vitelline membrane, which does not impair embryo growth, proliferation or mitotic apical constriction over experimentally relevant timeframes (Supplementary Figure 2A-D). DiI labelling, optimised to produce sparse mosaic labelling (Supplementary Figure 2E-F), enables individual cells to be tracked over time (Supplementary Figure 2G). Individual cells asynchronously displace their nuclei and either constrict or dilate during imaging (Figure 2C). We used nuclear morphology to identify cells which progressed through mitosis during live imaging, versus those that remained in interphase throughout (Figure 2D) and CellMask membrane stain to relate each nucleus or mitotic chromatin to its apical surface (Figure 2E). Live-imaged chick neuroepithelial cells undergo apical constriction in the 30 minutes preceding anaphase (Figure 2F). Both mitotic and interphase apical constrictions follow a pulsatile pattern (Figure 2G). To reconcile asynchronous changes, individual cell profiles were temporally aligned. T0 was defined as the timepoint when anaphase morphology was observed for mitotic cells and the smallest recorded apical area for non-mitotic cells. Individual cell traces were normalised to each cell’s average apical area to reveal trends independently of inter-cell variability (Figure 2G-H). Mitotic cells tend to dilate up to approximately 30 min prior to anaphase, then predictably constrict to a minimum apical area at anaphase before gradually re-dilating to an interphase-like pattern within 30 min following anaphase (analysing one daughter cell following cytokinesis) (Figure 2G). Following division, the daughter cells resume a pattern of lower-magnitude apical area fluctuations equivalent to those seen in cells which do not divide during live imaging. The pulsatile nature of individual cells’ apical constrictions can be approximated as a sine curve (Figure 2H). The constriction amplitude of mitotic cells is significantly greater than apical area oscillations observed in interphase (Figure 2I).

### Human iPSC-derived neuroepithelia apically constrict between G2 and M phase

Equivalent analyses are not possible in human embryos. Many protocols have been described to differentiate human iPSCs into neural progenitor cells, but few have been extensively characterised in the early stages of neuroepithelial induction to confirm transition through this morphogenetically-relevant stage. Additionally, most protocols induce a presumptive neuroepithelial stage as a cell aggregate with deep apical surfaces inaccessible to *en face* imaging. We sought to characterise a human system producing flat neuroepithelial sheets with accessible apical surfaces. Dual-SMAD inhibition (Shi et al., 2012b, Chambers et al., 2009) is well known to produce neurogenic cells in a flat sheet (Figure 3A), but its early differentiation stages are minimally characterised.

**Figure 3:**
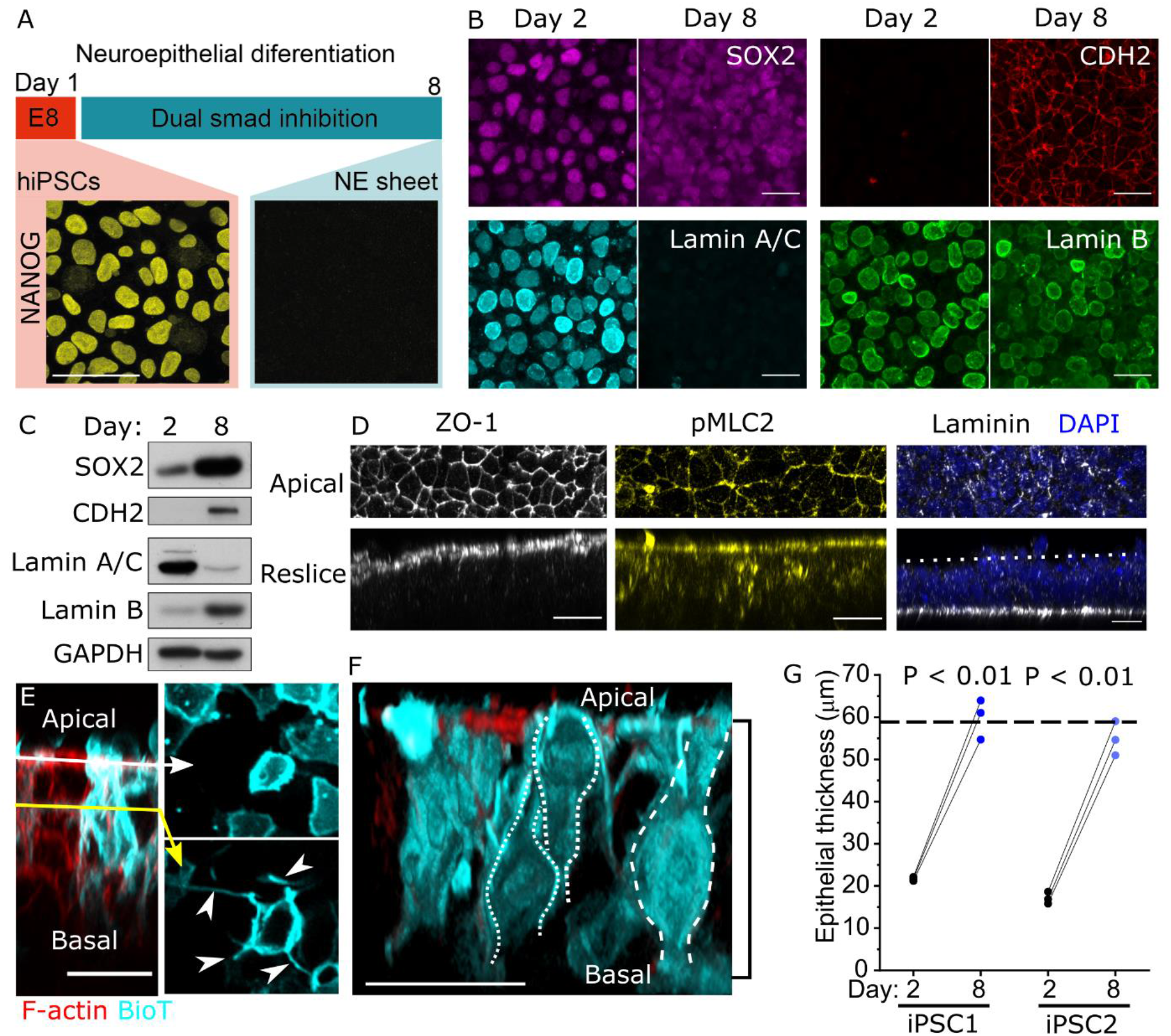
Characterisation of pseudostratified human iPSC-derived neuroepithelial sheets. **A.** Schematic of the 8-day differentiation protocol through dual SMAD inhibition. NANOG immunofluorescence is shown. Scale = 10 μm. **B.** Representative confocal images showing gain of CDH2, loss of Lamin A/C, and retention of SOX2 and Lamin B between days 2 and 8 of differentiation. Scale = 20 μm. **C.** Western blot visualisation of the markers in B. Note that as nuclear density increases between day 2 and 8, a greater proportion of the lysate may be nuclear when equal quantities of protein are loaded. **D.** Representative confocal images showing apical localisation of ZO-1, and phosphorylated (p)- MLC2, and basal localisation of laminin (dashed line = apical surface) in iPSC-derived neuroepithelia. Scale = 20 μm. **E.** iPSC-derived neuroepithelial cells stained mosaically with BioTracker (BioT). Apical (white arrow) and sub-apical (yellow arrow) optical sections are shown. Arrowheads indicate sub-apical protrusions. Scale = 20 μm. **F.** 3D reconstruction of BioT-trained iPSC-derived neuroepithelial cells. Dashed outlines indicate cell contours. The black bracket indicates epithelial thickness quantified in **G**. Scale = 25 μm. **G.** Quantification of neuroepithelial thickness on day 2 and day 8 of neuroepithelial differentiation. Points represent independent experiments, linked by black lines. P value by paired t-test. The dashed horizontal line indicates the typical neuroepithelial thickness of the mouse PNP (Galea et al., 2021).

Over 8-days of differentiation these cells lose pluripotency markers including NANOG (Figure 3A), but retain the dual iPSC and neuroepithelial marker SOX2 while gaining the neuroepithelium-selective adherens junction marker CDH2 (Figure 3B-C). A molecular peculiarity of neuroepithelial cells is their loss of Lamin A/C while retaining Lamin B around their nuclear envelope, both of which are recapitulated in these cultures (Figure 3B-C). This minimum panel of molecular markers defines a NANOG^-^/SOX2^+^/CHD2^+^/Lamin A/C^-^ epithelial population which, to our knowledge, is exclusive to neuroepithelial cells, complementing previously reported (Chambers et al., 2009, Shi et al., 2012b) extensive molecular characterisation of dual-SMAD differentiated cells.

Of greater consequence to the current studies is the morphology of the epithelial layer produced. Following eight days of differentiation, iPSC-derived neuroepithelial cells are apicobasally polarised with apical localisation of ZO-1 and enrichment of phospho-myosin, with a single continuous basement membrane containing laminin (Figure 3D). They form sub-apical lateral protrusions (Figure 3E) similar to those recently reported in vivo (Kasioulis et al., 2022). Each cell spans the apical to basal domain of the epithelium, which is as thick as the E9.5 mouse neuroepithelium (Figure 3F-G). Thus, this short protocol of human iPSC differentiation produces a tractable neuroepithelial sheet through which to mechanistically study cell behaviours such as apical constriction.

Both iPSC lines tested achieve apical areas equivalent to the mouse PNP (Figure 4A). Mitotic cells localise their rounded body apically, under a small apical surface (Figure 4B). In these human cells, Ki-67 can be used to segregate cells in different stages of the cell cycle as previously reported (Galea et al., 2013, Ghule et al., 2011). Ki-67 forms multiple nuclear foci in G1/S, coalesces into a single nuclear punctum in G2 and then spreads over the condensing chromosomes as cells enter M phase (Figure 4C). Mitotic neuroepithelial cells apically localise their nuclear chromatin below a ring of phosphorylated myosin (Figure 4D). Both human iPSC lines tested show variations of apical area of individual cells at each stage of the cell cycle, demonstrating that apical area variability is an inherent feature of neuroepithelia even in the absence of extrinsic chemical or mechanical inputs (Figure 4E). Both lines also show significantly larger apical areas in the G2 phase of the cell cycle, constricting to significantly smaller areas in M phase and retaining equivalent areas in G1/S (Figure 4E). Thus, G2 to M phase apical constriction is an intrinsic neuroepithelial behaviour conserved in human cells.

**Figure 4:**
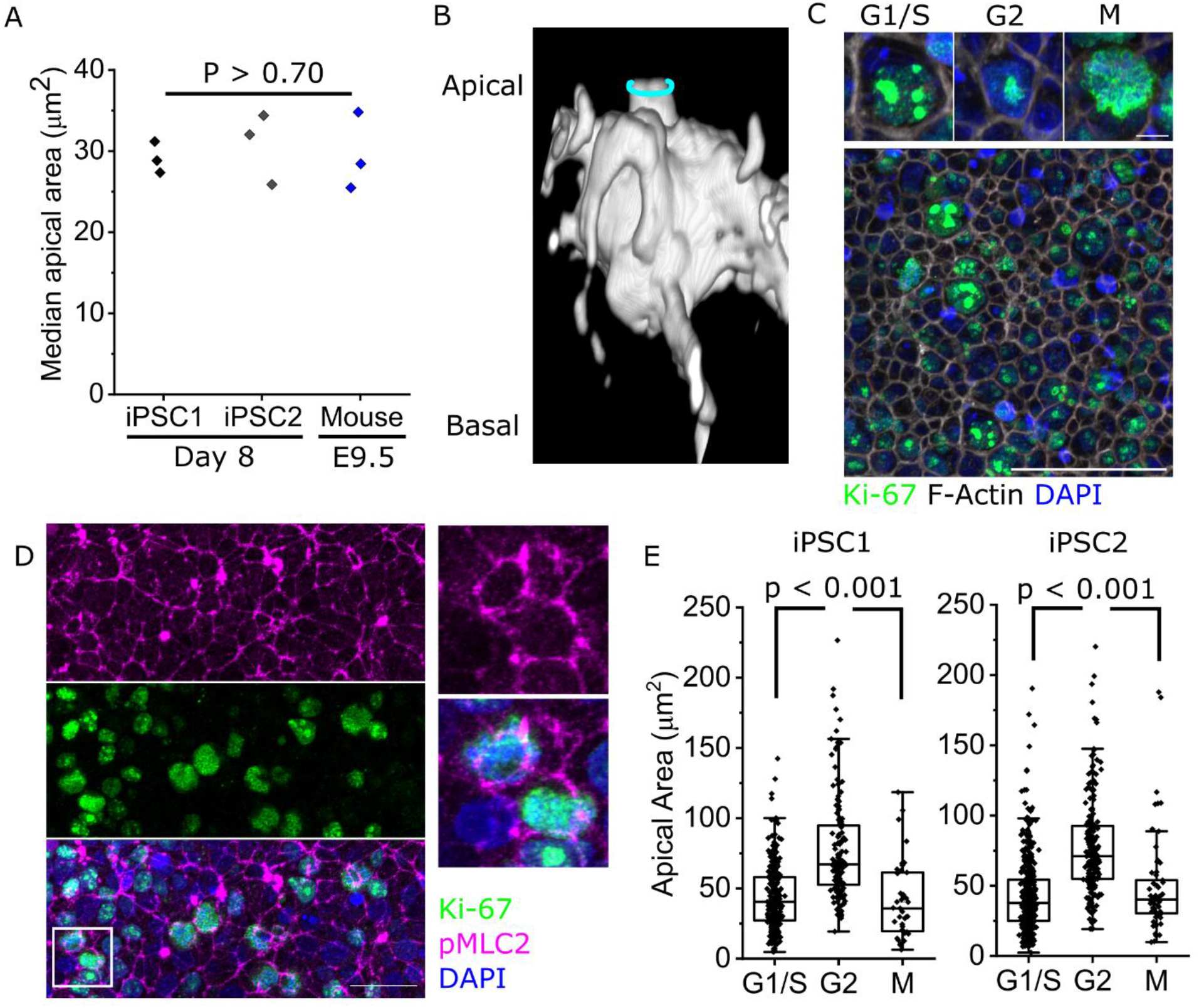
Human iPSC-derived neuroepithelia undergo mitotic apical constriction. All analyses are in cells after 8 days of differentiation. **A.** Comparison of neuroepithelial apical areas between the two human iPSC lines used in the current study after 8 days of differentiation, and 17-19 somite-stage mouse embryo PNP. Points represent independent experiments/embryos (note mouse values consistent with larger previous datasets). **B.** 3D reconstruction showing the shape of an illustrative BioT mosaically-labelled mitotic neuroepithelial cell. The cyan ring indicates the apical surface. **C.** Representative Ki67 staining panel showing identification of different cell cycle stages. Scale = 5 μm in inset, 50 μm in overall view. **D.** Representative confocal projection showing pMLC2 and Ki-67 double-labelling. The white box indicates the region shown in the inset. Scale = 25 μm. **C.** Apical area quantification of iPSC-derived neuroepithelial cells in G1/S, G2 or M phase. iPSC1: G1/S 283, G2 145, M 40 cells from 3 independent cultures. iPSC2: G1/S 502, G2 201, M 62 cells from 3 independent cultures. P value by ANOVA with post-hoc Bonferroni.

### Tissue geometry influences the timing of apical constriction during cell cycle progression

The demonstration of apical constriction in different species prompted us to test conservation of this behaviour in a different anatomical site, namely the anterior neuroepithelium of mouse embryos (Figure 5A). Contrary to the spinal region, anterior neuroepithelial cells in the presumptive midbrain do not undergo mitotic apical constriction, but rather dilate their apical surfaces, producing the largest apical areas observed in the epithelium (Figure 5B-D). MHC-IIb apical localisation shows both cortical and apical cap localisation in the midbrain region analysed (Figure 5E), as seen in the PNP. However, contrary to the PNP, no correlation is present between apical area and myosin staining intensity in either pHH3+ (Figure 5F) or pHH3- (Supplementary Figure 3A) anterior neuroepithelial cells. Consistent with this, ROCK is not notably enriched around the apical cortex of mitotic anterior neuroepithelial cells (Supplementary Figure 3B). Given that pHH3 labels cells from G2 to anaphase, we assessed whether the proportion of cells with corresponding nuclear morphologies differ between the anterior and spinal neuroepithelium, finding no differences (Supplementary Figure 4A). However, we observed that pHH3+ nuclei with a G2 morphology were on average closer to the neuroepithelial apical surface in the anterior than spinal neuroepithelium (Supplementary Figure 4B,C).

**Figure 5:**
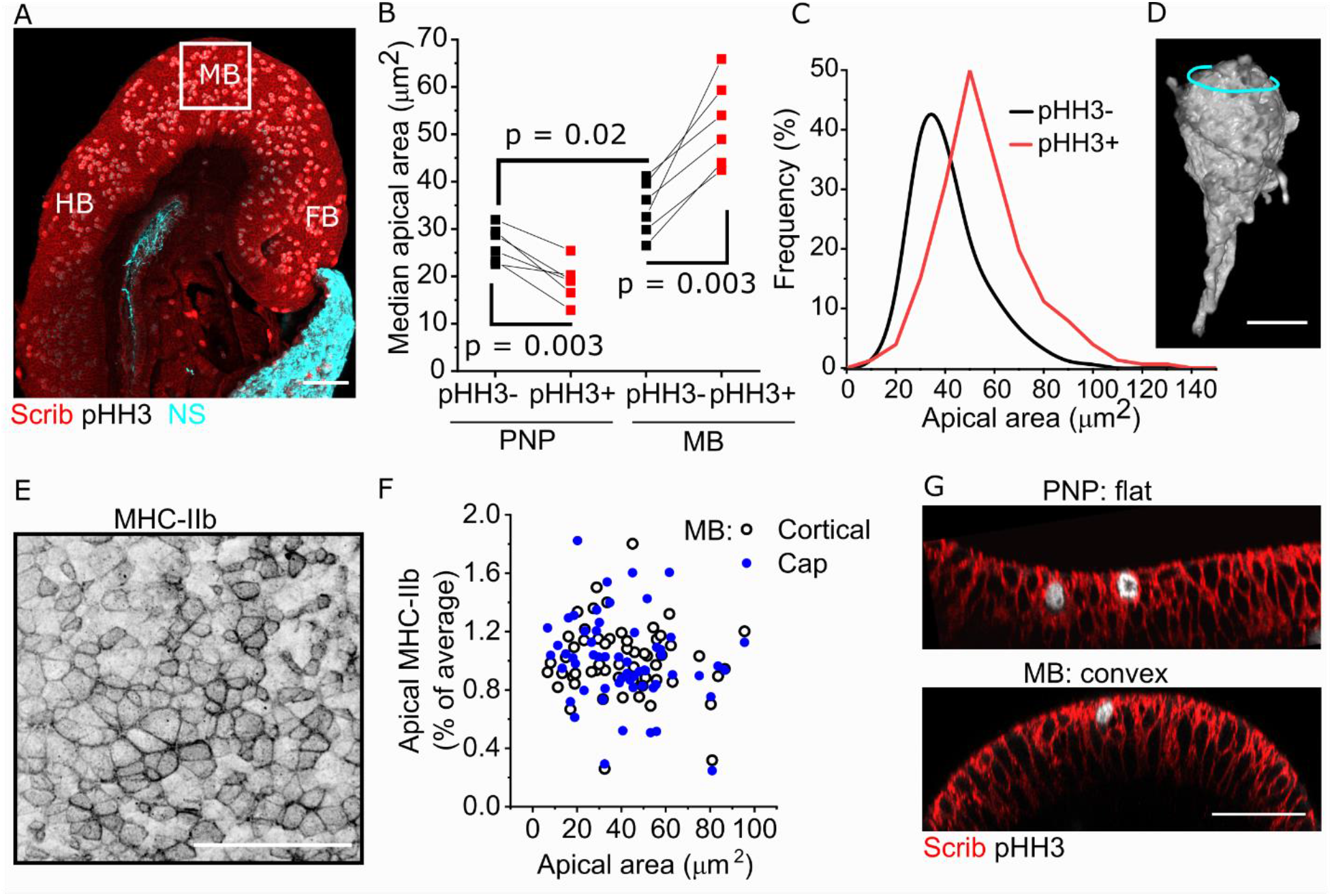
Mitotic apical constriction does not occur in the anterior neuroepithelium of the future mouse mid-brain. **A.** Medial wholemount image of a midline-sectioned 8-somite mouse embryo head indicating the fore- (FB), mid- (MB) and hindbrain (HB). The box indicates the region where analyses were performed (in dorsal-view images). NS non-specific (extraembryonic membranes). Scale = 100 μm. **B.** Quantification of median apical area in pHH30- versus pHH3+ cells in the PNP of E9.5 (19-22 somites) embryos and the midbrain of E8 (6-8 somite) embryos. Note the PNP apical surface is inaccessible to en face imaging at E8. Points represent individual embryos. P values from two-way ANOVA with post-hoc Bonferroni. **C.** Frequency plot of apical areas of pHH3+/- neuroepithelial cells in the early mouse midbrain. **D**. EGFP 3D reconstruction of a mosaically-labelled midbrain neuroepithelial cells. EGFP lineage-tracing was induced with Sox2^CreERT2^. Cyan line indicates the apical surface. Scale = 10 μm. **E.** Representative surface-subtracted MHC-IIb staining in a 7-somite embryo midbrain. Scale = 50 μm. **F.** Correlations between apical area and apical cap or cortical MHC-IIb intensity of pHH3+ cells. Each embryo’s MHC-IIb values are normalised to its average staining intensity to correct for inter-individual differences. Points represent 59 cells from four independent 6-8 somite embryos. **G.** Illustrative optically-resliced images showing the flat analysis region in the E9.5 PNP and apically convex region in the E8.5 midbrain. Scale = 50 μm.

Nuclear position in fixed embryos is a snapshot of their dynamic displacement apically. The molecular mechanisms and rate controlling this ascent have previously been shown to be influenced by neuroepithelial curvature (Yanakieva et al., 2019a, Ishii et al., 2021), which is substantially different between the relatively flat PNP versus the apically convex midbrain region analysed (Figure 5G). We therefore hypothesised that tissue geometry may determine whether neuroepithelial cells undergo mitotic apical constriction or dilation. Consistent with this, the median apical area of midbrain mitotic cells in individual embryos is significantly negatively correlated with their tissue’s radius of curvature, such that more acute apical curvature predicts larger mitotic apical areas (Supplementary Figure 4D,E).

To directly test the effect of geometry on mitotic apical constriction, we differentiated human iPSC-derived neuroepithelia on coated spherical glass beads mimicking early midbrain curvature (Figure 6A). Cells on the top of these beads did not form a pseudostratified epithelium and were excluded from further analyses, but those along the sides of the beads form a thick, apically convex neuroepithelium (Figure 6B,C). Their resulting apical curvature is variable, but on average is at least as curved as that observed in the early mouse midbrain (Figure 6D). Cells in these apically convex regions continue to apically localise cortical phospho-myosin (Figure 6E). As had been seen in flat cultures, the apical area of cells in G2 is significantly larger than those in G1/S in both iPSC lines tested (Figure 6F). However, M phase cells on curved geometries retain larger apical areas than in G1/S (Figure 6F). Thus, cell cycle-related regulation of apical area is an evolutionary conserved neuroepithelial behaviour, but mitotic apical constriction is selectively displayed by cells on flat tissue geometries.

**Figure 6:**
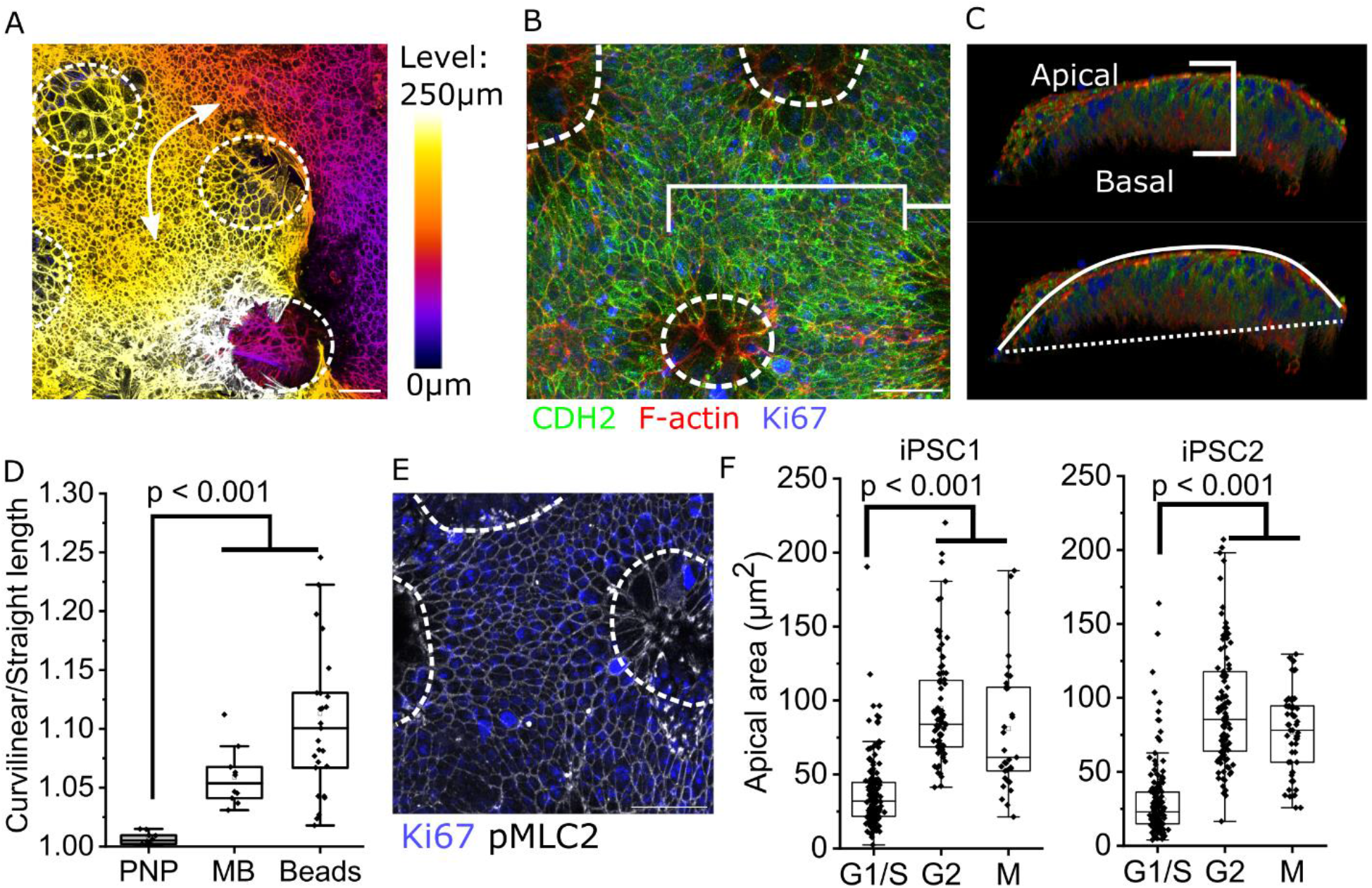
Mitotic apical constriction depends on epithelial geometry. **A.** Elevation projection showing the generation of curved epithelia (arrow) after eight days of neuroepithelial differentiation. Dashed circles indicate the position of beads used to generate curvature (throughout), which were excluded from analysis. Level indicates the elevation of the structures shown in maximum projection (0 μm = deepest level). Scale = 50 μm. **B.** Projected confocal image of a curved CDH2+ neuroepithelium after eight days of differentiation. Scale = 50 μm. **C.** 3D projection of B showing a thick, apicobasally polarised, pseudostratified, curved neuroepithelium. Curvature was quantified as the ratio between the curvilinear distance of the apical surface (solid line) to a straight connection (dashed line). **D.** Curvature calculation as in **C** comparing the PNP of E9.5 mouse embryos, the midbrain of E8.5 mouse embryos, and iPSC-derived neuroepithelia on curved beads. Points represent individual embryos or curved neuroepithelial cultures. P value from ANOVA with post-hoc Bonferroni. **E.** Projected confocal image of a curved neuroepithelium showing Ki67 and apical pMLC2 after eight days of differentiation. Scale = 50 μm. **F.** Quantification of apical area in each of the indicated cell cycle stages in curved neuroepithelia differentiated from two independent iPSC lines (three independent cultures each). P values from ANOVA with post-hoc Bonferroni.

## Discussion

Cells are often required to multitask, yet their molecular machinery is shared between functions. For example, cytoskeletal remodelling occurs during mitotic rounding and apical constriction, both of which can cooperate to drive morphogenesis (Kondo and Hayashi, 2013). Optimal balance between opposing behaviours is likely to depend on the wider context in which they occur, including the shape of their tissues. Here we show that mitotic apical constriction is a geometry-dependent behaviour which reverses apical expansion in flat neuroepithelia, but its absence prolongs dilatory effects of cell cycle progression in tissues with a convex apical geometry. Our findings during neurulation, when regulation of neuroepithelial apical size is a force-generating mechanism involved in closing the neural tube, complement studies of later developmental stages during which asymmetrical inheritance of the apical membrane and apical abscission contribute to neurogenesis (Nishizawa et al., 2007, Das and Storey, 2014).

In our previous report we documented that mitotic neuroepithelial cells in the mouse PNP have constricted apices relative to their interphase counterparts through mechanisms sensitive to pharmacological Rho/ROCK blockade (Butler et al., 2019). Here, we observe the highest apical non-muscle myosin levels in mitotic cells with the smallest apical surfaces, with good correlation between cortical and apicomedial cap myosin pools. This contrasts with invaginating *Drosophila* mesodermal and ectodermal cells in which apicomedial myosin decreases, and area increases, as cells enter mitosis (Ko et al., 2020). A difference between these *Drosophila* cells and mouse neuroepithelia is the abundance of cortical myosin in the latter. Cortical myosin predominates in *Vangl2-null* mouse PNP neuroepithelial cells which apically constrict more than their neighbours (Galea et al., 2021), suggesting this is the myosin pool primarily responsible for constriction. Consistent with this, junctional shortening is a well-established mechanism of neural plate apical constriction which follows pulsatile myosin accumulation during live imaging in *Xenopus* embryos (Ossipova et al., 2014).

Live imaging of the chick neuroepithelium shows that cell cycle stage is not the sole determinant of neuroepithelial apical area: the variability in apical dimensions of anaphase cells is larger than the average change during mitotic constriction. The apical surface area of the mouse anterior neuroepithelium decreases between E8.5 and E10.5 (after closure) (Ohmura et al., 2012, Grego-Bessa et al., 2016, Brooks et al., 2020), while retaining equivalent tension inferred from recoil following laser ablation (Bocanegra-Moreno et al., 2022). In the open PNP we observe that neuroepithelial apical areas are smaller in the chick than in the mouse, whereas apical areas of the mouse PNP and human iPSC-derived neuroepithelia are equivalent.

While other cultured epithelial cell types have been productively used to study apical constriction (Yano et al., 2021, Martin et al., 2019), to our knowledge, none are pseudostratified and undergo IKNM *in vitro*. Here, we characterise the robust differentiation of flat neuroepithelial sheets from human iPSCs through a previously-reported brief and simple dual SMAD inhibition protocol. The molecular identities of cells derived from this protocol have been extensively studied (Chambers et al., 2009, Strano et al., 2020), but we propose a minimum molecular markers panel which, to our knowledge, is unique to neuroepithelia. Of particular interest is this protocol’s relatively understudied ability to induce loss of the nuclear Lamin A/C, which facilitates IKNM in vivo (Yanakieva et al., 2019b). Several other iPSC differentiation protocols have been described which produce pseudostratified cells in 3D aggregates (Lee et al., 2022, Karzbrun et al., 2021, Veenvliet et al., 2021), but direct imaging of their apical surface would not be feasible in those systems. The heterogeneity of apical areas within individual cultures is striking, reflecting the in vivo situation despite lacking tissue-level morphology or external secreted cues. We are unable to comment on the dynamicity of their apical surfaces due to these cells’ apparent sensitivity to vital dye labelling and confocal/two-photon Z-stack imaging necessary to analyse their apical surface. Nonetheless, the granularity afforded by in situ cell cycle analysis using high-resolution imaging of human-specific Ki-67 (Galea et al., 2013) confirms that dilation occurs in G2.

IKNM nuclear displacement in G2 can be achieved by different molecular mechanisms depending on tissue geometry (Yanakieva et al., 2019a). In the flat zebrafish hindbrain, Rho/ROCK-dependent actomyosin activation rapidly displaces the nucleus apically, whereas in the concave retina sub-nuclear F-actin accumulation through formin polymerisation displaces the nucleus more gradually (Yanakieva et al., 2019a). In a static snapshot, slower ascent would produce a smaller proportion of apical nuclei. We observe more apical nuclei in the convex midbrain than the flat PNP of mouse embryos, consistent with the previous finding of slower nuclear displacement in concave than flat neuroepithelia (Yanakieva et al., 2019a). Differential rates of nuclear displacement during IKNM has also previously been live imaged in flat versus curved portions of the mouse cochlear epithelium (Ishii et al., 2021). One possible explanation for this is differential expression of actomyosin regulators, either linked to tissue-specific cell identities or secondary to cells’ responses to mechanical constraints.

Differential expression of relevant genes has previously been documented. For example, formin homology domain-containing (Fhod)3 is selectively expressed in apically convex regions of the anterior neuroepithelium overlying the rhombomeres and its deletion prevents apical constriction, causing exencephaly in mice (Sulistomo et al., 2019). In *Xenopus*, simultaneous accumulation of F-actin and N-cadherin (CDH2) parallels apical constriction anteriorly, whereas N-cadherin does not increase in the constricting posterior neuroepithelium (Baldwin et al., 2022). Important differences between *Xenopus* and amniotes include the lack of a pseudostratified neuroepithelium and brevity of neural tube closure relative to cell cycle progression in the former (Nikolopoulou et al., 2017). Given cell division progressively widens the PNP in the absence of apical constriction in mice (Butler et al., 2019), it is conceivable that synchronisation with cell cycle progression is of greater importance in slower neurulation events. Diminishing proliferation causes exencephaly, but rescues spina bifida in a mouse genetic model (Seller and Perkins, 1983, Copp et al., 1988). Opposition between cell cycle progression and PNP closure is consistent with computational analyses showing that IKNM, modelled without mitotic constriction, promotes apical expansion and nuclear crowding (Ferreira et al., 2019).

Apical expansion relative to the basal domain is characteristic of epithelia with convex geometries, such as the early mouse midbrain studied here. Subtle changes in tissue curvature change the effective force direction produced by cellular contractility (Maniou et al., 2021). Tissue curvature, and the challenges it poses for epithelial cells arrangements, is emerging as a topic of active research (Lou et al., 2022, Prabhakara et al., 2022, Gómez-Gálvez et al., 2022). Differential cell packing and stress peaks may explain the unexpected differentiation of non-neuroepithelial, squamous cells at the top of glass beads used in these studies. The identity of these CDH2-negative cells is unknown, yet they remain connected to the pseudostratified cells which differentiate along the curved sides of each bead. These cells do show dynamic cell cycle-related changes in apical area: a clear increase between G1/S and G2 as well as constriction between M and G1/S. However, their imposed geometry abolishes mitotic apical constriction. This may be an adaptive process which facilitates expansion of the apical surface relative to the basal, or reflect a limitation of mitotic cells ability to mechanically pull their neighbours’ cell junctions over their apical rounded body.

Thus, synchronization between the cell cycle and apical area is a conserved feature of neuroepithelia, potentially counteracting apical expansion promoted by IKNM (Ferreira et al., 2019). Failure of this synchronization may impair neural tube closure by causing progressive widening of the neuropore as cells divide (Butler et al., 2019). This synchronization is conserved in a tractable and validated human iPSC-derived model. The ability to empirically change these cells’ geometry corroborates in vivo observations that mitotic apical constriction does not happen in apically convex neuroepithelia, retaining larger apical areas until they return to the G1 phase. Persistent dilation during M phase correlates with the expanded apical relative to basal surface in convex epithelia. These findings suggest neuroepithelial cells are not only able to multitask by balancing growth through proliferation with neuropore shrinkage through apical constriction, but do so while interpreting the physical cues of their tissue structure.

## Materials and Methods

### Mouse embryo collection

Studies were performed under project and personal licenses regulated by the UK Animals (Scientific Procedures) Act 1986 and the Medical Research Council’s Responsibility in the Use of Animals for Medical Research. Pregnant female mice were time-mated and sacrificed by cervical dislocation 8.5 or 9.5 days after the morning a plug was identified. C57Bl/6J mice were bred in-house and mated when at least six weeks old. mTmG reporter mice used to visualize cell outlines (Muzumdar et al., 2007), Nkx1.2^CreERT2^ used to mosaically label PNP cells (Rodrigo Albors et al., 2018), Sox2^CreERT2^ used to mosaically label anterior neuroepithelial cells (Andoniadou et al., 2013) bred in house, and their administration of oral tamoxifen (50 μl of 100 mg/ml solution in corn oil per mouse), were all as previously described (Savery et al., 2020, Galea et al., 2021).

### Chick embryo culture and live imaging

Fertilised Dekalb white chicken eggs (Henry Stewart, Norfolk, UK) were incubated at 37°C in a humidified chamber for ~36 hrs reaching Hamburger and Hamilton stages 9-10. Embryo collection and culture was according to the EC protocol (Chapman et al., 2001) with the slight modification of utilising a double filter paper sandwich as a carrier. Any excess yolk was washed off with Pannett-Compton saline. Vitelline membrane windowing was done with a tungsten needle.

Embryos were first stained with a 1:1 mix of 1:50 Hoechst (ThermoFisher Scientific #62249) and 3:10,000 ice-cold DiI suspension (Invitrogen #D282) in PBS for 15 min at 37°C. Following washing with Pannett-Compton saline, embryos were further stained with 1:100 CellMask Deep Red plasma membrane stain (Invitrogen #C10046) in PBS at 37°C for 15 min. Any excess stain was washed off with Pannett-Compton saline. Live imaging was done with a Zeiss Examiner LSM880 confocal under AiryScan Fast mode using a 20x/NA0.7 C-Epiplan Apochromat dry objective. Hoechst was excited using a MaiTai tuneable two-photon laser to avoid phototoxicity.

### Human iPSC culture and neuroepithelial differentiation

The previously established human hiPSC lines were used: SFC086-03-03 (iPSC1) was obtained from EbiSC bank and HO-193b (iPSC2) was as previously published (Michielin et al., 2020). For the culture of hiPSCs in E8 medium (Thermo Fisher Scientific), cells were maintained on 0.5% Matrigel® Growth Factor Reduced (GFR) (Corning 354230) coated tissue culture plates, as described (Michielin et al., 2020, Beers et al., 2012). When confluency reached ~80% cell colonies were washed with PBS without CaCl_2_/MgCl_2_ (Gibco) and dissociated with 0.5 mM EDTA (Thermo Fisher Scientific) for 3-6 minutes at 37°C. Cells were resuspended in E8 medium and re-plated at a 1:6 split ratio. All cell lines were regularly tested for mycoplasma contamination.

The protocol for direct differentiation of hiPSCs into neuroepithelial sheets was as previously described (Shi et al., 2012a), with minor modifications. Cells differentiated following this protocol can form neurons with a dorsal forebrain identity (Shi et al., 2012a). Briefly, cells were lifted using EDTA solution and plated in 1% Matrigel®-coated tissue culture plates. Approximately ~280,000 cells/cm^2^ were transferred into a 35 mm glass bottom dish (ibidi, Thistle Scientific) ensuring 100% confluency the following day. After 24 hours, the cells were washed twice with PBS and medium was replaced with neural induction medium consisting of an equal ratio of N2 (N2 supplement in DMEM/F12, Thermo Fisher 17502048 and 31331093) and B27 solution (B27 supplement in Neurobasal, Thermo Fisher 17504044 and 12348017) supplemented with 10 μM SB431542 and 1 μM Dorsomorphin (both Tocris Bioscience). The medium was changed daily, and cells were cultured for up to eight days.

### Immunofluorescent staining, confocal microscopy and image analysis

Embryo were stained for a minimum of 4 hours in cold 4% paraformaldehyde (PFA) and wholemount staining as previously described (Galea et al., 2017). Primary antibodies used were mouse anti-phospho-histone H3 (S10, Cell Signalling Technology antibody #9701), rabbit anti-ROCK (Abcam antibody ab45171), rabbit anti-Ser19 pMLC (Cell Signalling Technology antibody #3671), rabbit anti-total MHC-IIb (Abcam antibody ab230823), mouse anti-NANOG (Abcam antibody ab173368), rabbit anti-SOX2 (Merck antibody ab5603), mouse anti-CDH2 (Cell Signaling Technology antibody #14215), mouse anti-Lamin A/C (SantaCruz antibody sc7292), goat anti-Lamin B1 (SantaCruz antibody sc6216), rabbit anti-Ki67 (Abcam antibody ab16667), rabbit anti-ZO1 (Thermo Fisher Scientific antibody #40-2200), rabbit anti-laminin (Abcam antibody ab11575) and detected with Alexa Fluor™ conjugated secondary antibodies (Thermo Fisher Scientific). ROCK immunolabelling required prior antigen retrieval in 10 mM, pH 6, sodium citrate solution for 25 minutes at 90°C. F-actin was labelled with Alexa Fluor™-647 conjugated phalloidin (Thermo Fisher Scientific). Images were captured on a Zeiss Examiner LSM880 confocal using a 20x/NA1 or 10x/NA0.5 Plan Apochromat water immersion objectives. AiryScan Opt or Flex settings were used with automatic processing in ZEN. Stereoscope images were captured using a Leica DFC490 camera mounted on a Zeiss Stemi SV-11 stereomicroscope.

Images were processed and visualized as 3D or maximum projections in Fiji (Schindelin et al., 2012). PureDenoise and 5-pass Richardson Lucy deconvolution in Deconvolution Lab (Sage et al., 2017) were applied to live-imaged chick datasets (distributed by the Biomedical Imaging Group, EPFL). The elevation map was generated using the Fiji temporal LUT, replacing T with Z. Surface subtraction was performed essentially as previously described (Galea et al., 2018) using an in house Fiji macro available from https://github.com/DaleMoulding/Fiji-Macros (courtesy of Dr Dale Moulding).

Cell apical areas were manually segmented in 3D confocal Z-stacks, following the cell’s outline from its nucleus to its apical surface. The actomyosin cortex is readily identifiable as a bright rim around each cell’s apical surface. The inner border of this cortex was used to define an area selection representing the apical cap, in which myosin intensity was quantified. The perimeter of this area selection was then converted into a line and expanded away from the cell’s centre by 0.5 μm. The resulting band was used to define the apical cortex. Thicknesses, intensity, length, and areas were measured using in-built Fiji functions. In each case, every mitotic cell in each field of view was analysed and at least as many non-mitotic cells were analysed in the same field of view, analysing contiguous non-mitotic cells whenever possible.

Mosaic labelling of neuroepithelial cells was achieved with BioTracker™ 490 Green Cytoplasmic Membrane Dye (Thermo Fisher Scientific). Fixed cells were stained according to the manufacturer’s instructions for 30 minutes at room temperature.

### Western blotting

Western blotting was performed essentially as previously described (Thompson et al., 2022, Galea et al., 2020). Cells were lysed on ice for 30 min in 50 mm Tris Base pH 7.6, 150 mm NaCL, 1% Triton X-100, 0.02% Sodium Azide, 1 mm protease inhibitor cocktail (Merck Life Science UK Ltd), 1 mm sodium orthovanadate, 25 mm sodium fluoride. Protein concentration was determined using Pierce BCA Protein Assay kits (Thermo Fisher). Proteins were resolved by SDS-PAGE using 10% polyacrylamide gels and transferred to PVDF membranes. Membranes were blocked in TBS (15.4 mm Trizma-HCL, 4.62 mm Tris-base, 150 mm NaCl, pH 7.6) containing 10% w/v milk powder (Merck Life Science UK Ltd). Membranes were incubated with primary antibody overnight and then with horseradish peroxidase (HRP)-conjugated secondary antibodies (Agilent Technologies, Stockport, UK) for 1 h. Primary antibodies were as described in the immunofluorescence section. HRP was detected using ECL Prime (Cytiva, Amersham, UK). To strip and re-probe, membranes were incubated in 0.2 m NaOH for 20 min at 37°C and then 20 min at room temperature before re-blocking and re-use.

### Statistical analysis

Comparison of two groups was by two-tailed t-test, paired when measures were repeated (e.g. same cell at two time points). Comparison of more than two groups was by one-way or two-way ANOVA with post-hoc Bonferroni correction. Linear regression was by Pearson’s correlations. Sine curve fitting and all statistical tests were performed in Origin 2021. P < 0.05 was considered statistically significant. Blinding was not possible due to obvious differences in tissue geometry or the need to identify individual cells.

Wherever possible, the embryo or independent culture dish (cultured on different days) was considered the unit of measure. When this was not possible, e.g. due to small numbers of mitotic cells in each not providing a representative value, the cell was considered the unit of measure. All mitotic cells fully present in an imaged field of view were included in each analysis. All representative data is based on at least four independent (different culture/litter) observations.

## Author contributions

GLG designed the study in discussion with EM, PDC, NE, AJC, NDEG and FL. IA performed all iPSC work and analysis with help from EMT, TW, FP, GGG, and CM. GLG performed all mouse work and analysis with help from CE and EM. Chick embryo work and analysis was performed by CE with help from EM and GLG. GLG, EM and PDC supervised students. GLG, PDC, NE, AJC, NDEG, FL, CM, GGG and FP provided resources. GLG drafted the manuscript, and all authors approve the version submitted.

## Acknowledgements

This study was supported by the Wellcome Trust (211112/Z/18/Z and 211112/Z/18/A, both to GLG), the Child Health Research CIO (20-21/SI2) and an NIHR Great Ormond Street Hospital Biomedical Research Centre PhD studentship to IA. The views expressed are those of the authors and not necessarily those of the NHS, the NIHR or the Department of Health.

