## Supplementary for "Synchronisation of apical constriction and cell cycle progression is a conserved behaviour of pseudostratified neuroepithelia informed by their tissue geometry"

### Supplementary Figures and Legends:

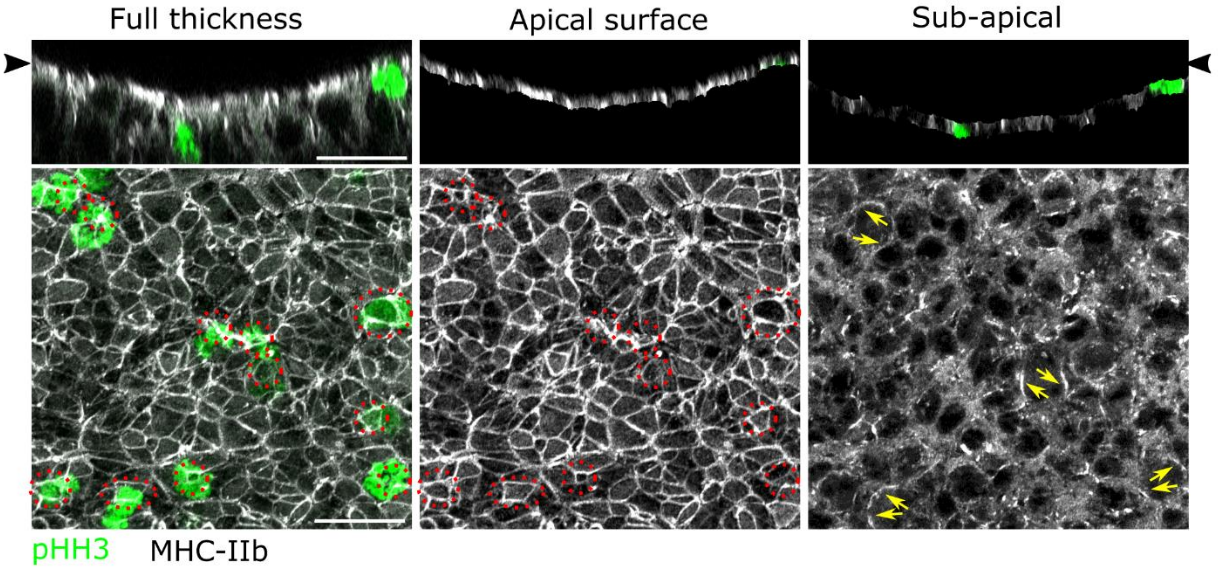


**Supplementary Figure 1: Mitotically rounded neuroepithelial cells retain apical MHC-IIb.** *Serially surface-subtracted wholemount image of a 17-somite mouse embryo PNP showing apical (dashed rings) and sub-apical (yellow arrows) myosin localisations. Arrow heads indicate the apical surface in optically resliced images. Scale = 25 µm.*

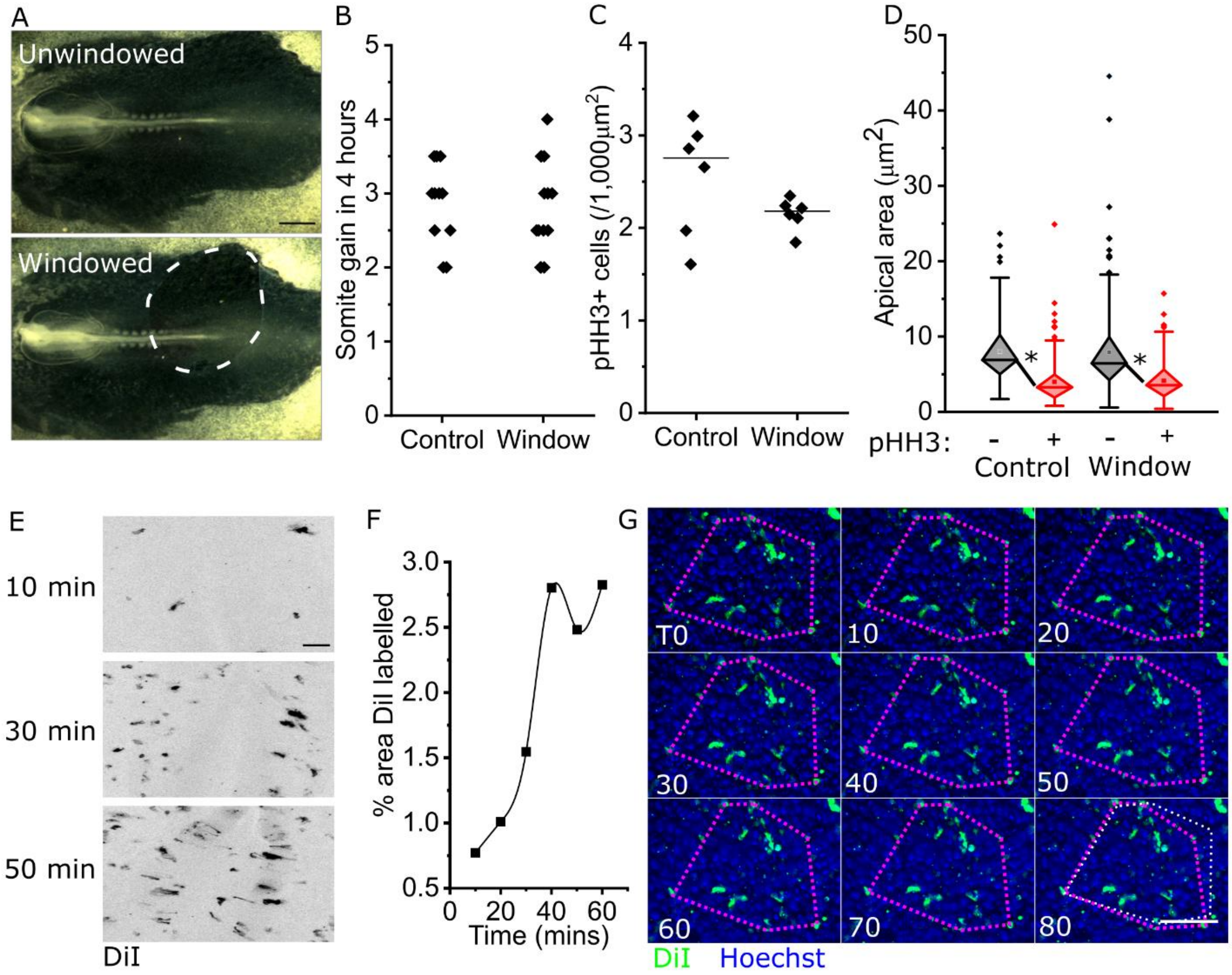


**Supplementary Figure 2: Vitelline membrane "windowing" does not impact embryo growth or mitotic apical constriction.**

*Chick embryos in EC culture had a portion of the vitelline membrane removed and cultured for a further four hours before fixation. Control embryos were placed in culture at the same time but not subjected to windowing.*

***A.*** *Illustrative brightfield images of an embryo before and after windowing the vitelline membrane (dashed line). Scale = 100 µm.*

***B.*** *Quantification of the number of somites gained over 4 hours in control and windowed embryos.*

***C.*** *Comparison on mitotic rate between control and windowed embryos.*

***D.*** *Comparison of apical area between mitotic and non-mitotic cells in control and windowed embryos. * $p < 0.001$ by two-way ANOVA with post-hoc Bonferroni (no differences between control and windowed samples). Control = 176 pHH3+ and 176 pHH- cells from 4 embryos (distinct from Figure 2B). Windowed = 211 pHH3+ and 211 pHH3- cells from 5 embryos.*

***E-F.*** *Optimization of chick posterior neuropore staining with DiI.* ***E)*** *Fluorescent stereoscope images showing the increase in DiI labelling in embryos incubated with the staining solution for the indicated lengths of time. Shown in inverted grey LUT (black = DiI-stained cells), scale = 50*

*µm.* ***F)*** *Images of DiI-stained PNPs were binarized and the proportion of area labelled quantified. Each dot represents an individual embryo.*
***G.*** *Sequential live-imaging of a chick PNP mosaically labelled with DiI at the indicated time points (minutes). The magenta dashed polygon at each timepoint identifies the same cells located at its vertices. The polygon changes shape as the cells it interconnects change position. The white dashed polygon at T = 80 minutes indicates the original shape at T0. Scale = 50 µm.*

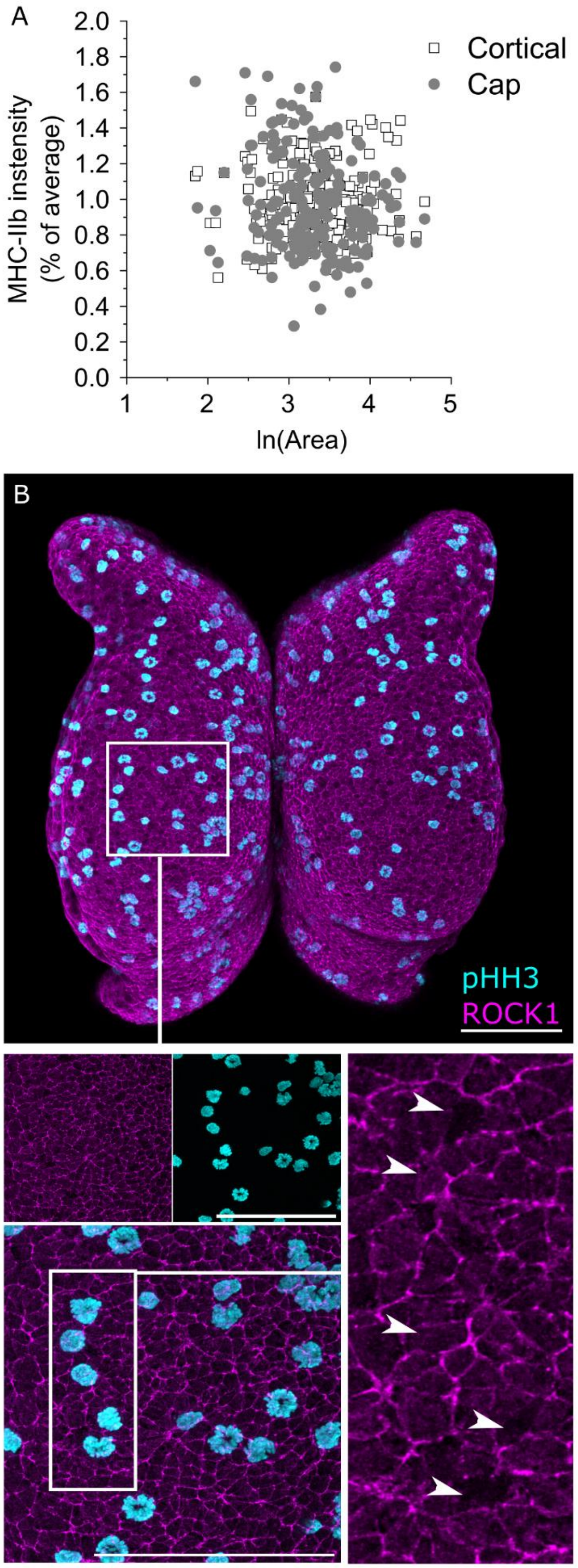


**Supplementary Figure 3: Mitotic anterior neuroepithelial cells do not show selective apical ROCK localisation.**
***A.*** *Lack of correlation between apical area and cap or cortical MHC-IIb staining intensity in pHH3- anterior neuroepithelial cells. Each embryo's MHC-IIb values are normalised to its average staining intensity to correct for inter-individual differences.*
***B.*** *Wholemount dorsal-view of a 7-somite stage embryo. Arrowheads indicate the apical surfaces of mitotic cells. Scale = 100 µm.*

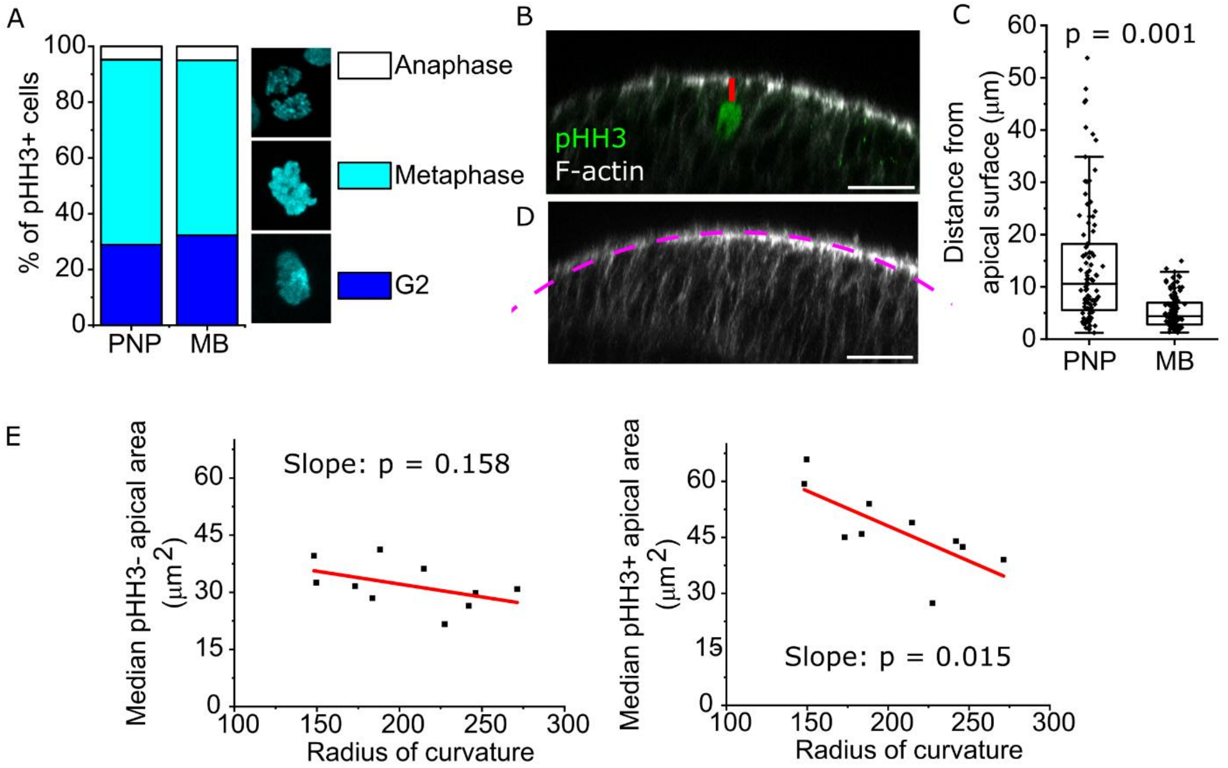


**Supplementary Figure 4: Influences of geometric inputs on neuroepithelial characteristics.**
***A.*** *Proportion of pHH3+ nuclei with G2, metaphase (brightest signal) or anaphase morphology in the PNP of E9.5 (19-22 somites, 211 cells from six embryos) embryos and the midbrain of E8 (6-8 somite, 175 nuclei from five embryos) embryos.*
***B.*** *Optical cross-section through a 6-somite stage midbrain. The red line indicates the distance between a pHH3+ cell with a G2 nuclear morphology and the apical surface. Scale = 25 µm.*
***C.*** *Quantification of the distance between G2 nuclei and the apical surface in the E9.5 PNP (96 cells from 12 embryos) and E8.5 midbrain (109 cells from 11 embryos). P value by T-test accounting for significant difference in variance (F-test $p < 0.001$).*
***D.*** *Illustration of a circular arc (dashed magenta line) fitted to calculate the radius of curvature of the apical midbrain neuroepithelium. Scale = 25 µm.*
***E.*** *Correlation between the midbrain radius of curvature (smaller radius = more acutely curved) and the median apical area of pHH3+ and – cells in that embryo. Points represent individual 6-9 somite stage embryos. Slope p values from Pearson's correlation.*
